# Climate-fire-vegetation interactions and the rise of novel landscape patterns in subalpine ecosystems, Colorado

**DOI:** 10.1101/290163

**Authors:** W. John Calder, Ivanka Stefanova, Bryan Shuman

## Abstract

1. Feedbacks at multiple scales can be important for shaping how forest ecosystems respond to both climate change and disturbance. At landscape scales, feedbacks likely exist between vegetation and wildfire regimes such that a change in one produces changes in the other. More locally, some forest patterns can result from feedbacks between plants and their abiotic environment. Alternating areas of forest and meadow (ribbon forests) in subalpine forest provide an example where both scales of feedbacks could be important with changes in climate-vegetation-fire interactions giving rise to local-scale feedbacks between snow drifting and forest extent that created the ribbon forests and further feedback to alter fuel continuity and fires regimes.
2. To examine the feedbacks in subalpine forests and the history of ribbon forests, we obtained six fossil pollen records from lakes across a subalpine landscape in Colorado. Forests there may have responded to climate change and widespread wildfires ca. 1000 years ago when >80% of sites on the landscape burned within a century. The fires coincided with regional warming, but the extent of burning declined before the climate cooled, possibly driven by changes in fuel structure and composition.
3. Results of cluster analyses of the pollen percentages indicate that large changes between successive sets of samples coincided with the widespread wildfires at five of the six sites. After the wildfires, sagebrush (*Artemisia*) and other meadow taxa increased as conifers, especially spruce (*Picea*), declined across the landscape, indicating that the forests opened.
4. *Synthesis.* The opening of the forests may have created fuel breaks across the mountain range that limited wildfire after temperatures rose ~0.5 °C. When the openings then became larger and the area covered by ribbon forest expanded during the Little Ice Age (LIA), the extent of fires further declined. Pollen assemblages associated with modern ribbon forests only became common across our study sites during the LIA when the frequency of fires across our sites reached its minimum. The rise of novel ribbon forests in northern Colorado thus illustrates how climate and fire can interact to rapidly transform landscapes and their disturbance regimes.

## Introduction

Recent increases in the area burned per year by wildfire and other forest declines around the world challenge ecologists to consider how disturbances may permanently alter vegetation patterns and composition (Allen et al., 2010; Turner, Gardner, & O’Neill, 2015). Disturbances may catalyze large-scale ecological responses to climate change, which could include novel ecological communities and patterns (Jackson, Betancourt, Booth, & Gray, 2009; Millar & Stephenson, 2015; Turner, 2010). By creating persistent stress, climate change can alter the potential responses to disturbance, and thus enable disturbance events to trigger critical transitions to new ecosystem states (Scheffer, Carpenter, et al., 2012). Such changes may feedback to further alter the disturbance regimes, and thus permanently alter ecosystem function as well as structure and composition, and paleoecology can provide retrospective insights on such dynamics (e.g., Clifford and Booth 2015).

Here, we consider how climate, wildfire, and vegetation interacted in a subalpine ecosystem over the past two millennia. The setting is ideal for examining of how climate changes and disturbances interact with each other and with vegetation patterns for several reasons. First, subalpine forests support stand-replacing fires, which leave a sedimentary record because of the accumulation of charcoal in lakes. The fires, in turn, influence the composition and structure of the forests, which is recorded by fossil pollen. Second, high-elevation forests grow at their climatic limits and are, thus, sensitive to changes in climate over time. Finally, climate can determine the frequency of wildfires, which often burn most extensively during warm, dry years. Subalpine ecosystems, thus, exemplify the triangular relationship among climate, fire regimes, and vegetation (Fig. 1; Whitlock et al. 2010).

**Fig. 1.**
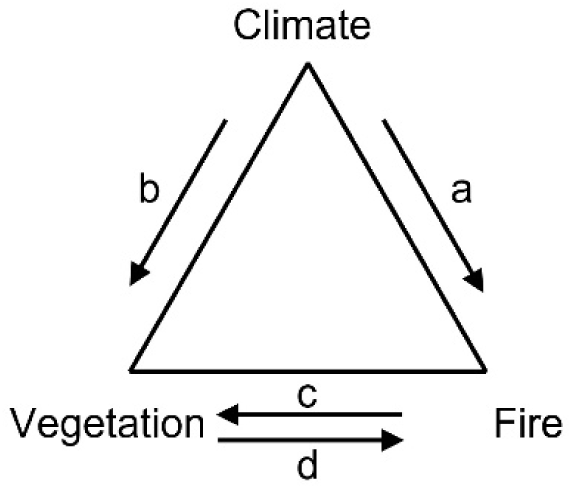
Fire regime triangle indicating the relationship between climate change, wildfire, and vegetation. Climate influences both vegetation and wildfire, and vegetation and wildfire can interact to influence each other.

We ask whether the interactions among climate, fire, and vegetation can produce landscape-scale state shifts by examining fossil pollen records across a mountainous landscape where both climate and fire regimes changed substantially in recent millennia (Fig. 2). Our previous work in this ecosystem showed that the area burned per century was sensitive to temperature changes with more than 80% of our study landscape burning when mean annual temperatures rose ~0.5 °C during Medieval times about 1000 years ago (Calder, Parker, Stopka, Jiménez-Moreno, & Shuman, 2015). However, the high rates of burning did not persist as long as the region remained warm, suggesting ecological limits to sustained, large wildfires. We hypothesized that warming facilitated larger fires, but that the fires drove changes in vegetation structure that limited the spread of additional fires (Calder et al., 2015). Consistent with this hypothesis, we also found that the fires at one of our study sites accelerated the local development of “ribbon forests”, a discontinuous mix of linear alpine meadows and ribbons of conifer forest (Fig. 3; Calder & Shuman, 2017). Fossil pollen indicates that ribbon forests only developed there after the fires, but were maintained throughout the last millennium by climatic cooling, which culminated in the Little Ice Age (LIA), a period of cool conditions beginning about AD 1300 or 650 BP (years before AD 1950)(Calder & Shuman, 2017).

**Fig. 2.**
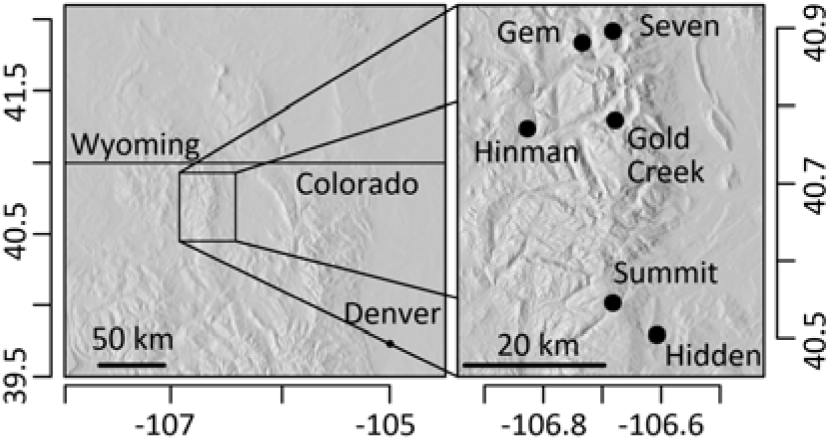
Coring locations for the lake sites in and around the Mount Zirkel Wilderness Area in northern Colorado.

**Fig. 3.**
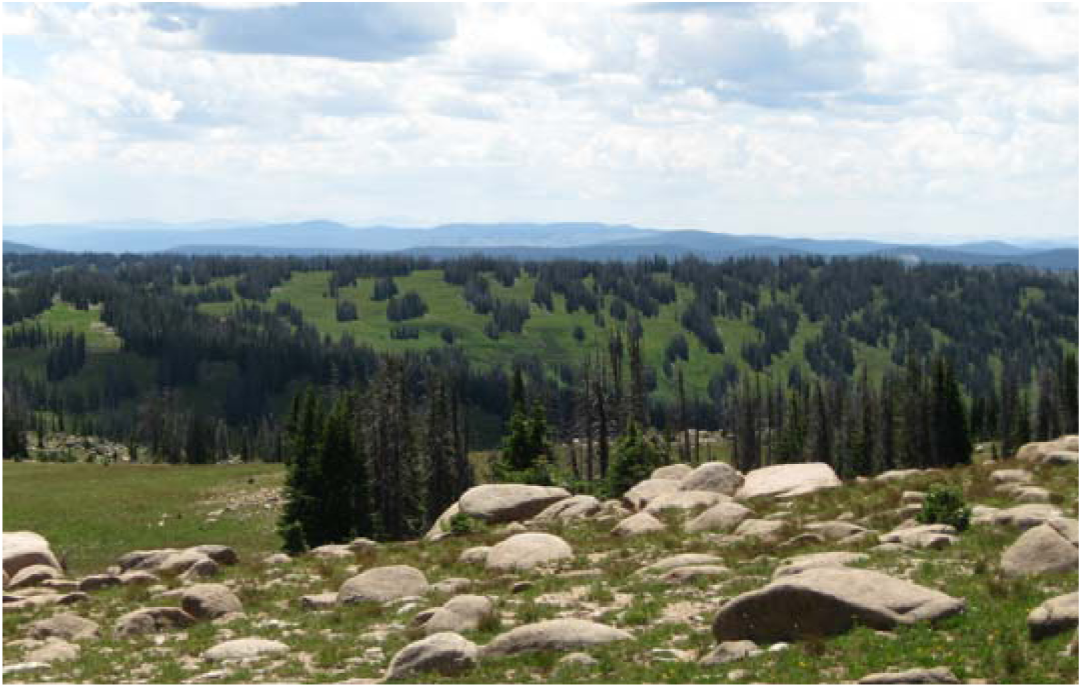
Ribbon forests within our study area.

Using additional records, this paper further evaluates two paired hypotheses about the interactions of climate, fire, and vegetation in subalpine ecosystems. We hypothesize that large fires can 1) trigger persistent legacies in the pattern of vegetation across a landscape when climate trends prevent a return to pre-fire states, and 2) produce vegetation changes that feed back to alter fire regimes. Potentially, such interactions can give rise to novel vegetation patterns, which may be the case here if ribbon forests had not previously existed within this landscape.

To examine these hypotheses, we reconstructed the vegetation history of the landscape that includes the Mount Zirkel Wilderness of northern Colorado (Fig. 2). We generated five new fossil pollen records from across a range of elevations (Table 1). Including our previous study site (Calder & Shuman, 2017), the network of records includes six lakes with two lakes within discontinuous ribbon forests, two lakes within mid-elevation conifer forests adjacent to areas of ribbon forest, and two other lakes within low elevation conifer-aspen forests. All of these locations show evidence of the large fires at ca. 1100 years before AD 1950 (BP) (Calder et al., 2015), and can thus provide constraints on the extent and persistence of rapid post-fire vegetation changes.

**Table 1.**
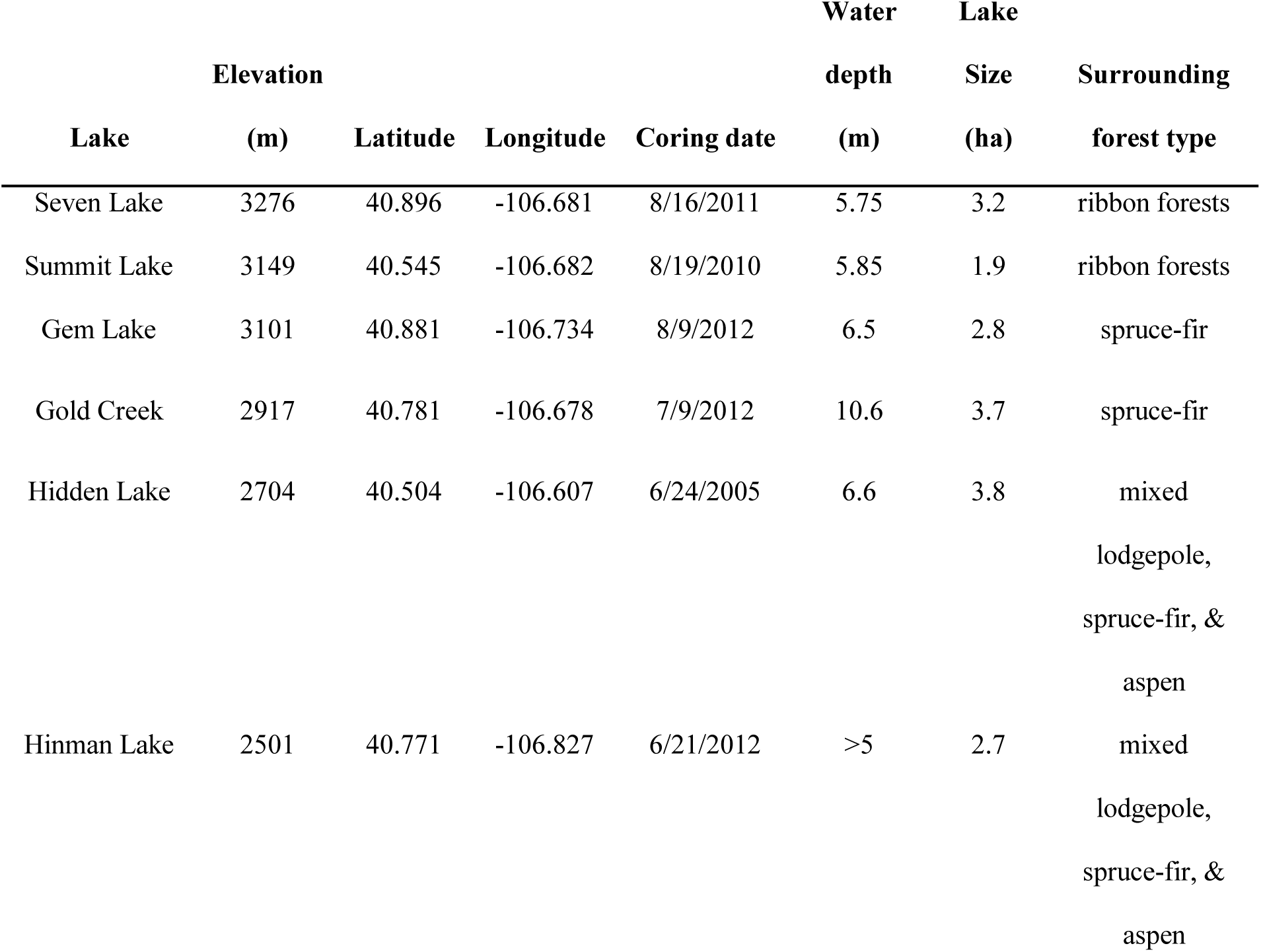
Lake site information

## Study Sites

Our study area spans across the Park Range, which forms the northern most range along the Continental Divide in Colorado and is located primarily within the Mount Zirkel Wilderness area of the Routt National Forest. Each of the lakes cored are similar in size, between 1.9 and 3.8 ha, and similar in depth, between 5 and 10 m deep (Table 1). Elevations within the study area range from 2000 – 3700 m with greater topographic relief and variability in the northern than southern half of the range (Fig. 2). Crystalline bedrock, including quartz monzonite, felsic gneiss, and mica schist dominates the geology (Snyder 1980a,b), and the range was also heavily glaciated (Atwood, 1937).

Across the lowest elevations, between approximately 2000 and 2800 m, mixed *Pinus contorta* (lodgepole pine) and *Populus tremuloides* (aspen) forests dominate the forest vegetation with some *Picea engelmannii* (Engelmann spruce) and *Abies lasiocarpa* (subalpine fir), particularly on north-facing slopes and along drainages. On the west side of the divide, these mixed forests adjoin a zone of *Quercus gambelii* (Gambel’s oak) with some stands of oak intermixing within the mosaic of mixed forest on south-facing aspects. Above approximately 2800 m, the vegetation is dominated by spruce-fir forests. At the highest portions of the mountain range, approximately >3100 m, the spruce-fir forests transition to ribbon forests and open meadows with patches of spruce-fir forests. The ribbon forests are composed of alternating bands of spruce-fir forests approximately 10 – 20 m wide and separated by 30 – 70 m wide meadows (Fig. 3; Billings 1969). Snowdrifts tend to persist late into the summer in these meadows and may play a role in the pattern and formation of the ribbon forests (Hiemstra, Liston, & Reiners, 2002; Moir, Rochelle, & Schoettle, 1999). The open meadows between the ribbons of spruce-fir forests are dominated by grasses, such as *Danthonia intermedia* and *Deschampsia cespitosa*, *Artemisia scopulorum* (sagebrush), and flowering plants in the *Asteraceae* family.

The two high elevation lake sites, Seven Lakes and Summit Lake, lie near the Continental Divide and ribbon forests currently surround both the lakes (Table 1; Calder et al. 2014). Summit Lake lies along a broad plateau with large areas of ribbon forests (Calder et al., 2014). By contrast, Seven Lakes lies along a narrow ridge where ribbon forests grow but cover only a small fraction of the source area for the pollen deposited on the lake, assuming a pollen source area of ~6 km radius (Schwartz, 1989).

Ribbon forests also lie within the pollen source areas of two other mid elevation sites, Gold Creek and Gem lakes. (Table 1). The primary vegetation surrounding Gem Lake is closed spruce-fir forests, but ribbon forests and open meadows lie 200 m upslope, within 300 m laterally, of the lake. Similarly, closed spruce-fir forests form the primary vegetation surrounding the north-facing slopes around Gold Creek Lake, which sits on the south side of a glacially-carved canyon. Ribbon forests and a mix of open meadows cover much of the nearby landscape within 1 km where elevations rise 300 m above the lake. At these high elevation lakes (Gem, Gold Creek, Summit and Seven Lakes), lodgepole pine and other pine species are rare in the areas surrounding the lakes, which indicates that any *Pinus* pollen found at these spruce-fir sites likely comes from long-distance transport.

Finally, mosaics of spruce-fir, lodgepole pine, and aspen forests surround the two lowest elevation sites, Hidden and Hinman lakes. West of the continental divide, open sagebrush parks and dense Gamble’s oak thickets, combined with spruce-fir, lodgepole, and aspen forests, grow at the low-elevation forest ecotone near Hinman Lake (Table 1). On the east of the divide, Hidden Lake is surrounded primarily by lodgepole pine and spruce-fir forests, with aspen groves further upslope.

## Methods

The work presented here builds on previous analyses of sediment cores collected from the six lakes. Calibrated radiocarbon chronologies and sedimentary charcoal stratigraphies used here are the same as previously published (Calder et al., 2015). Wildfire events, indicated by peaks in the rate of charcoal accumulation within each core, were calculated using standard techniques and form the basis for a reconstruction of the percentage of sites burned per century across the landscape (Calder et al., 2015). To generate the wildfire reconstructions, charcoal counts from contiguous 1 cm intervals (1-2 cm^3^ of sediment) were decomposed to identify fire episodes (Higuera, Brubaker, Anderson, Hu, & Brown, 2009). The age uncertainty distributions of the individual fire episodes were calculated using Bchron that models sediment accumulation rates using a Monte Carlo Markov Chain simulation in R (Haslett & Parnell, 2008; Parnell, Haslett, Allen, Buck, & Huntley, 2008; R Development Core Team, 2014). By combining the age uncertainty distributions for all fire episodes across all sites, including six additional charcoal records not used here for pollen analyses, we estimated the potential range of study sites burned per century for the last 2000 years (Calder et al., 2015).

### Pollen preparation and analyses

To detect the effects of the past fires on vegetation, we added fossil pollen analyses to five of the six sediment cores. A separate paper describes the detailed pollen record from the sixth core collected from Summit Lake (Calder & Shuman, 2017). In all cases, pollen was processed with standard techniques of acid digestion (Faegri, Kaland, & Kzywinski, 1989), and a minimum of 300 terrestrial grains per sample were counted. Pollen percentages were calculated from the sum of terrestrial taxa, excluding aquatic and wetland taxa, such as *Cyperaceae*. *Pinus* percentages were the combined counts from Diploxylon and Haploxylon types. Two researchers counted the pollen (Calder and Stefanova), and we tested for and corrected researcher-specific biases in the counts.

In addition to evaluating changes in the percentages of pollen from individual taxa, we considered changes in the ratio of pollen from the dominant trees (conifers) versus the taxa representative of open meadows. The ratio of conifer pollen (*Pinus*, *Picea*, and *Abies*) to the dominant subalpine non-arboreal taxa (*Artemisia*, *Poaceae*, and *Asteraceae*) was calculated using the terrestrial percentages. For simplicity, we refer to these groups as the conifers and herbs and shrubs, and refer to the ratio between them as the conifer:herb pollen ratio (C:H) to avoid confusion with the conventional arboreal:non-arboreal pollen ratio used in many palynological studies. Low-elevation taxa found primarily in the surrounding intermountain basins (e.g., *Sarcobatus*) were excluded. Lynch (1996) found that the ratio of C:H can discriminate between pollen assemblages from closed subalpine forests and high elevation treeless parks.

At the landscape scale, which we define as the scale represented by all six study sites together, we calculated the median C:H from all of the mid- and high-elevation sites (Gem Lake, Gold Creek Lake, Seven Lakes, and Summit Lake). One challenge making comparisons between pollen records arises from the age uncertainty associated with the individual sediment samples from each lake. To account for the age uncertainty between lakes, we linearly interpolated each pollen record 2250 times to 65-year time steps (the median pollen sampling resolution across sites) using 2250 different age-depth relationships generated in Bchron (Haslett & Parnell, 2008; Parnell et al., 2008). We then calculated the mean time series of the ratios for all lakes, where each 65-year time step had a C:H ratio averaged across all four sites from each 2250 different interpolation possibilities. We then use the median of the ensemble of four lake means, as well as 95^th^ and 5^th^ percentiles of the distribution, for our analysis. For the mean C:H from each lake record, we evaluated the probabilities of a change point at every time step using Bayesian change point analysis with the *bcp* R package (Erdman & Emerson, 2007)

Constrained cluster analysis (Grimm, 1987) was also used to evaluate changes in pollen assemblages within each pollen records using taxa with > 2% representation and clusters constrained to include only temporally adjacent sets of samples using the R package *rioja* (Juggins, 2015). The chi-square dissimilarity metric was used to create the dissimilarity matrix for each cluster analysis with the *analogue* R package (Simpson, 2007) because previous analyses showed the chi-square dissimilarity metric offered the best separation of dissimilarity among forests types across our study area (Calder & Shuman, 2017). We compare the timing of the large fires within the study area to the timing of breaks between clusters with the expectation that, if fires altered the composition of the vegetation, the timing of fires and cluster breaks should be correlated.

Unconstrained cluster analysis (not constrained by stratigraphic position) was calculated with the fossil samples from all six pollen records in the same matrix without age information. To do so, we used the chi-square dissimilarity and the same taxa list as the constrained cluster analysis. Previous work using a network of pollen surface samples from the area showed that pollen samples from the vegetation types of open forests near treeline and closed forests below treeline were distinguishable from ribbon forests (Calder & Shuman, 2017). Therefore, we used the first three clusters from the unconstrained cluster analysis to determine the distribution of the major vegetation types through time across the network of sites.

## Results

### Pollen Percentages

Elevational differences between the pollen source areas of each lake appear to affect the pollen assemblages (Figs 4 – 6). Summit Lake (Fig. 4b), which is functionally the highest site as the broad plateau allows for the greatest amount of ribbon forests, contains high *Picea* (>20%) and *Artemisia* (>40%) and low *Pinus* (<40%) pollen percentages. The records from the sites near large areas of mid-elevation forests (Seven, Gem, and Gold Creek lakes) contain intermediate percentages of *Pinus*, *Picea*, and *Artemisia* pollen. The pollen percentages from Seven, Gem and Gold Creek lakes fall between the high *Picea* and *Artemisia* and low *Pinus* percentages at Summit Lake and the low *Picea* (<10%) and *Artemisia* (<20%) and high *Pinus* (>50%) pollen percentages of the low-elevation lakes, Hidden and Hinman lakes (Figs 4 – 6). Hidden and Hinman lakes also contain the highest percentages (~5%) of *Quercus* pollen (Fig. 6).

**Fig. 4.**
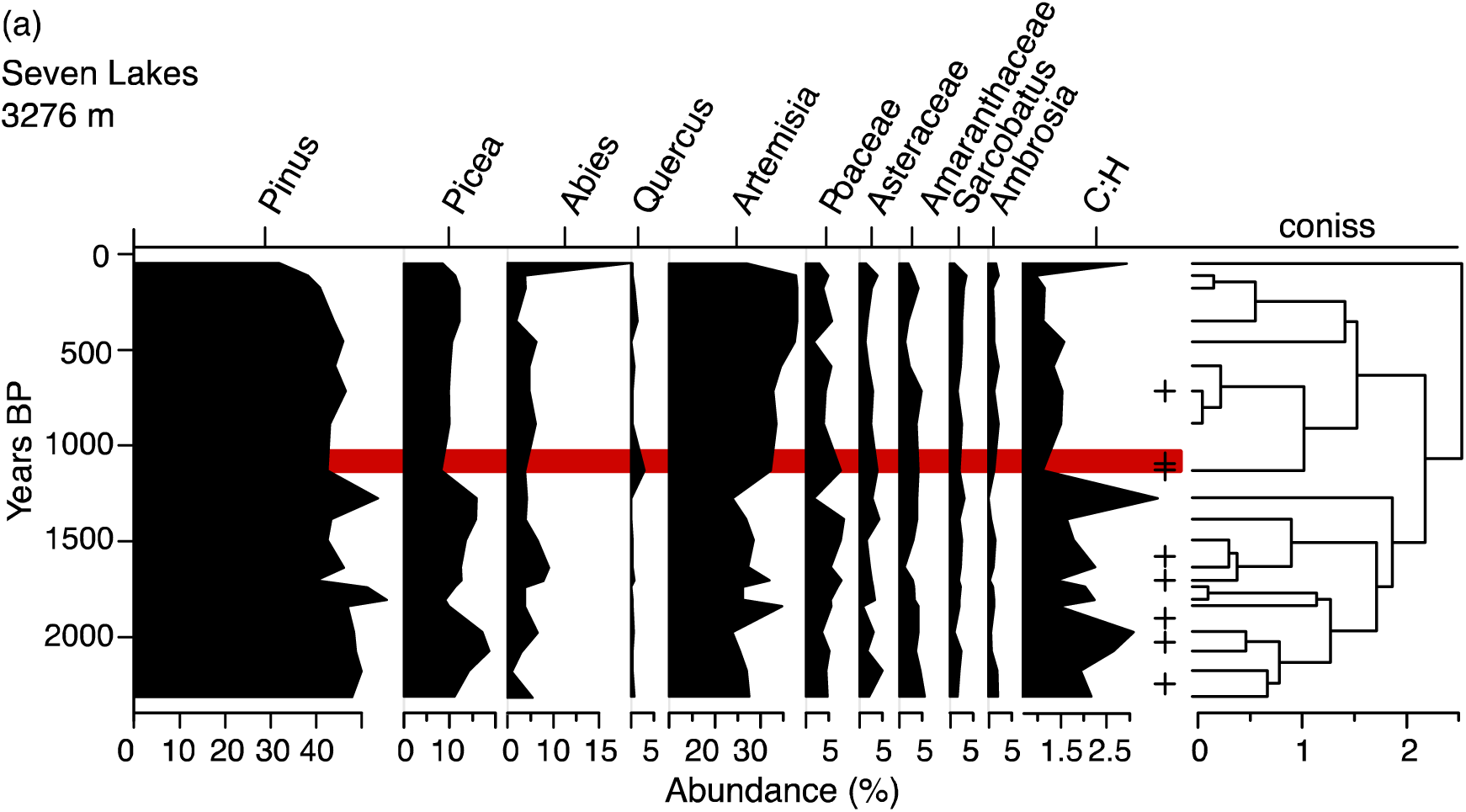
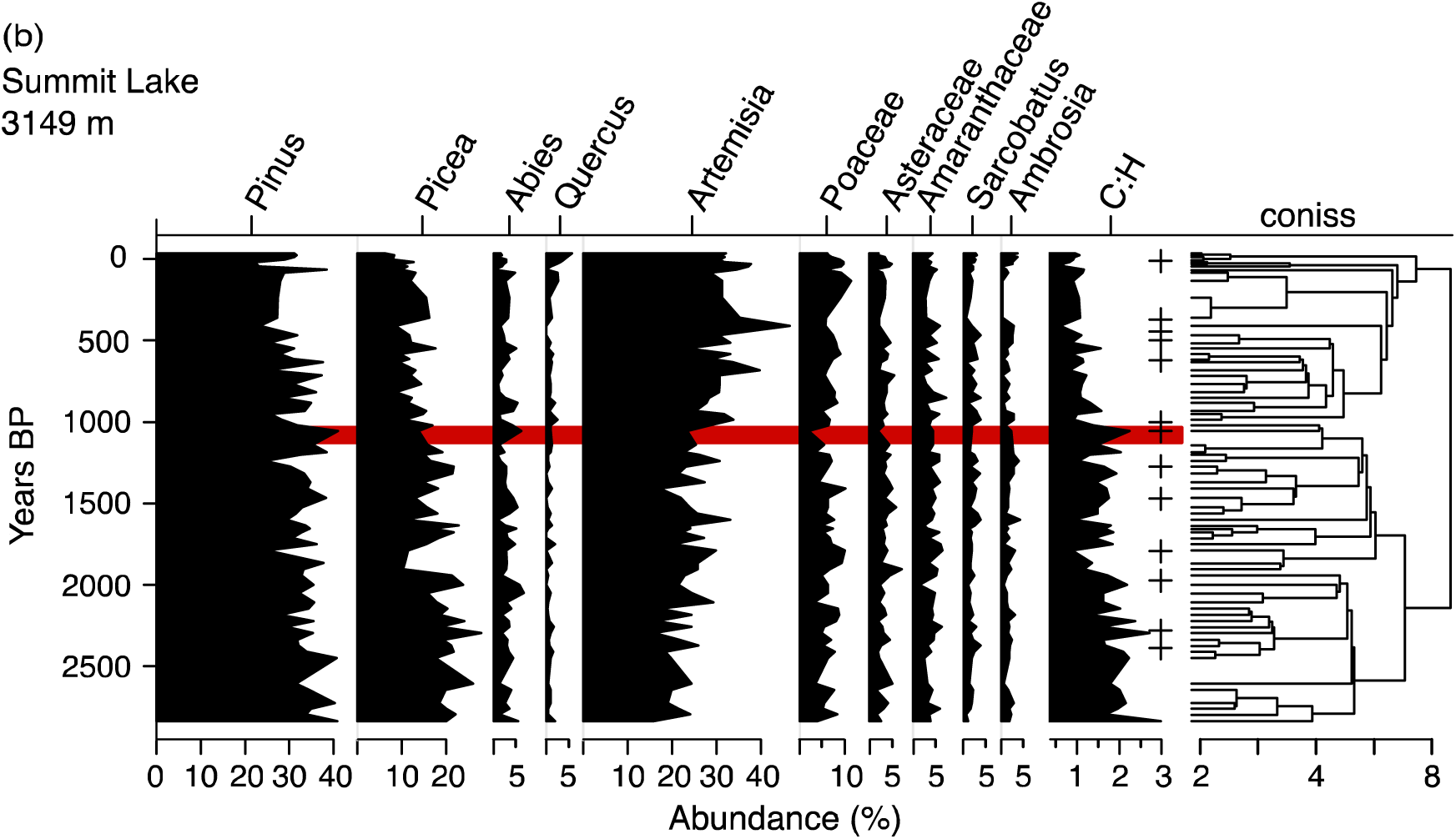
Terrestrial pollen percentages from the highest elevation sites. Plus symbols represent fire events detected within the individual cores and red bars highlight the century of peak landscape scale wildfires shown in Figure 7.

In most of the pollen records, however, changes in *Pinus*, *Picea*, and *Artemisia* pollen percentages define the important differences with time, with the minor taxa (*Sarcobatus, Ambrosia,* and Amaranthaceae) differing little through time (Figs 4 – 6). *Pinus* pollen percentages declined towards the top of each core at all sites, except for Hidden Lake, and *Artemisia* pollen percentages increased as *Pinus* declined. At Hidden Lake, *Pinus* and *Artemisia* percentages varied through time, but with no consistent trend (Fig. 6a). Several of the sites, particularly Seven and Gold Creek lakes, also contain peaks in *Quercus* pollen percentages at ca. 1000 BP and again in the last 300 years (Figs 4a and 5b).

At Summit Lake, *Picea* pollen percentages declined from >25% to <10% in the last 1000 years, especially after a sharp rise in *Artemisia* pollen percentages to >30% at 987 BP associated with the local expression of the large fires (plus symbols within the red bar, Fig. 4b). The maximum *Pinus* pollen percentages were reached just prior to this transition. The decline in *Pinus* at 1021– 987 BP coincides with a step shift in C:H from >2 to <1 (Fig. 4b) and a 0.77 probability of a change point at 1035 BP in the interpolated C:H.

At Seven Lakes, important changes also include a sharp decline in *Picea* pollen percentages to <10% from >15% by 1132 BP when two charcoal peaks represent the local expression of the extensive fires (plus symbols within the red bar, Fig. 4a). As at Summit Lake, a brief peak in *Pinus* pollen percentages, which reached >50%, preceded the decline at Seven Lakes (Fig. 4). The changes produced a shift in the mean C:H from 1.5 to 1 that distinguishes the periods before and after ca. 1000 BP and a 0.98 change point probability beginning at 1235 BP in the interpolated C:H.

Changes at Gem Lake include a decline in *Pinus* pollen percentages from >40% to ~30% at ca. 1000 BP in association with the charcoal peaks representative of the large fires (plus symbols within the red bar, Fig. 5a). Like the highest elevation sites, Summit Lake and Seven Lakes, *Pinus* pollen percentages at Gem Lake reached their maximum just before that decline. *Artemisia* rose subsequently and obtained its local maximum after 780 BP.

**Fig. 5.**
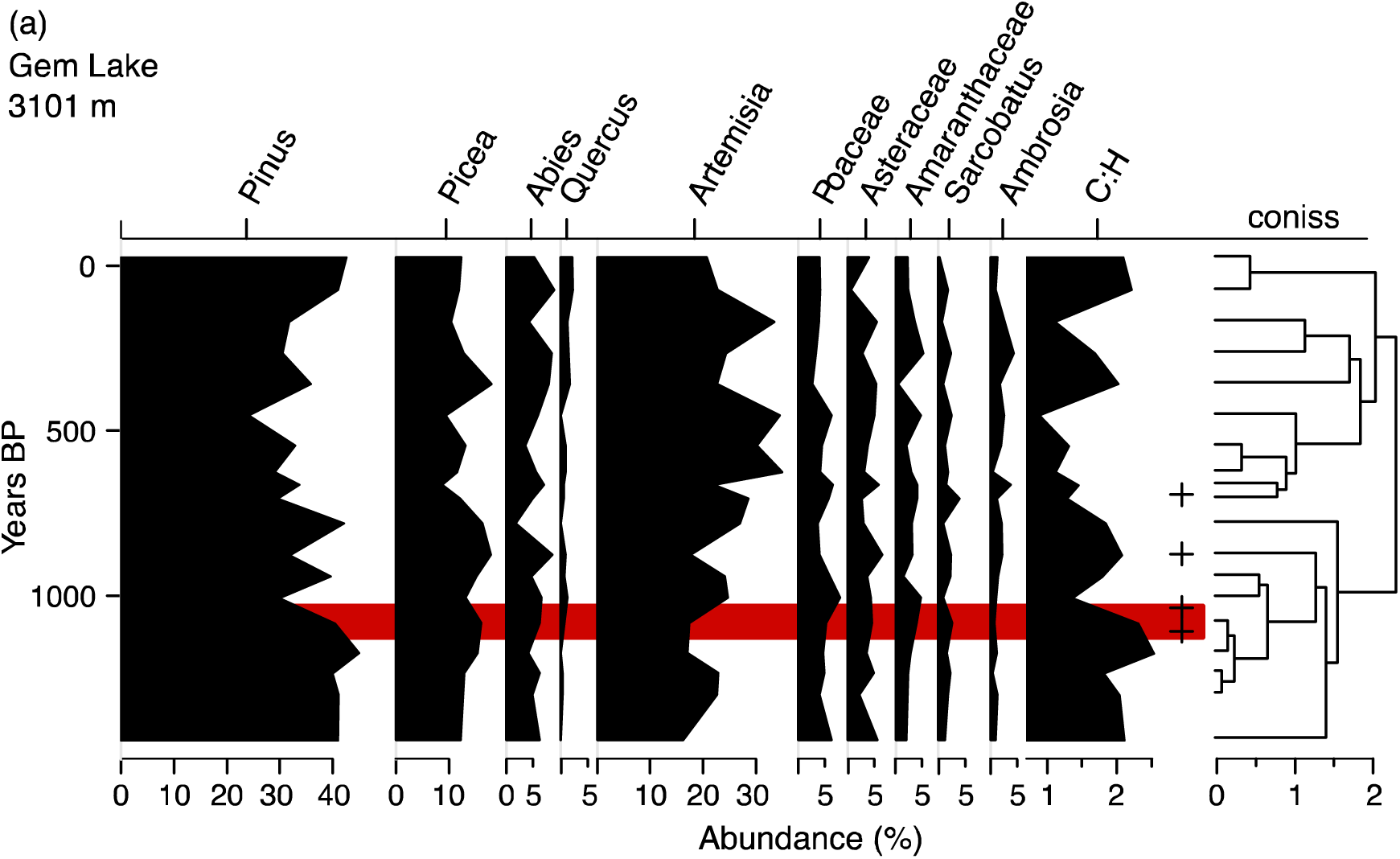
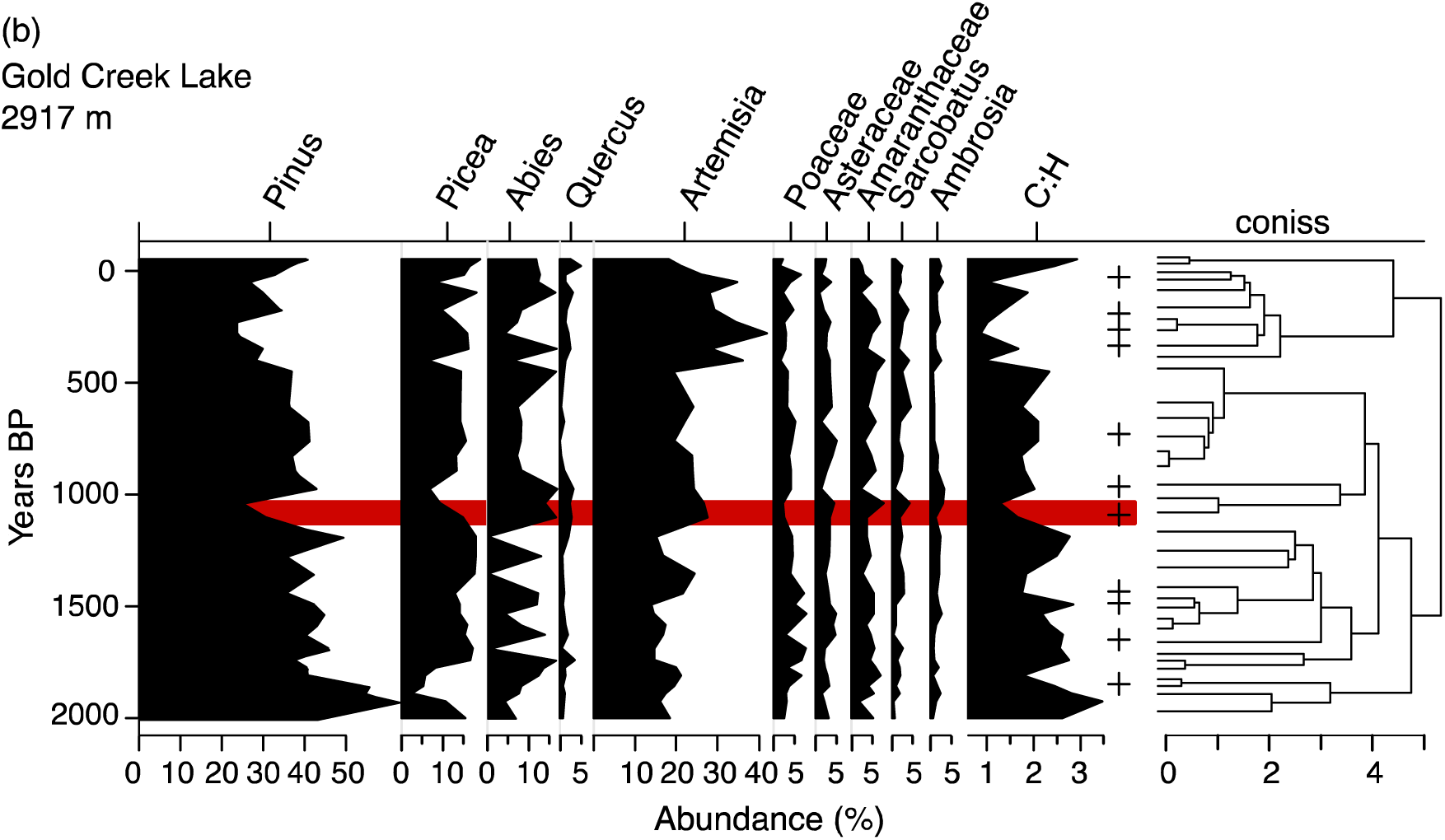
Terrestrial pollen percentages from the mid elevation sites. Plus symbols represent fire events detected within the individual cores and red bars highlight the century of peak landscape scale wildfires shown in Figure 7.

At Gold Creek Lake, *Pinus* and *Picea* pollen percentages declined sharply at 1101 BP in association with the local expression of the widespread fires, and when *Artemisia* pollen percentages increased stepwise by >10% (Fig. 5b). The C:H ratio also declined to <2 after 1101 BP and remained below 2 for the rest of the record. At 1165 the interpolated C:H declines with a 0.65 probability of change point. *Artemisia* pollen percentages did not reach their local maximum (>30%), however, until after another sharp >10% increase after 500 BP. An early phase of high *Pinus* pollen percentages, like those observed at Hidden and Hinman lakes (>50%), dates to 2000 – 1800 BP at Gold Creek, but another short-lived maximum preceded the fires at 1188 BP.

The pollen record from Hidden Lake contains more variability in C:H ratios than the other lakes with a sharp decline at ca. 1500 BP and a sharp increase at ca. 500 BP (Fig. 6a). The changes represent a tradeoff between high *Pinus* and *Artemisia* pollen percentages, which culminated in a *Pinus* minimum and *Artemisia* maximum associated with peaks in Poaceae and herbaceous taxa at ca. 600 BP after the only fire episode detected from 1000 – 100 BP. The interval at ca. 600 BP includes the highest rates of charcoal accumulation in the core (Calder et al., 2015).

**Fig. 6.**
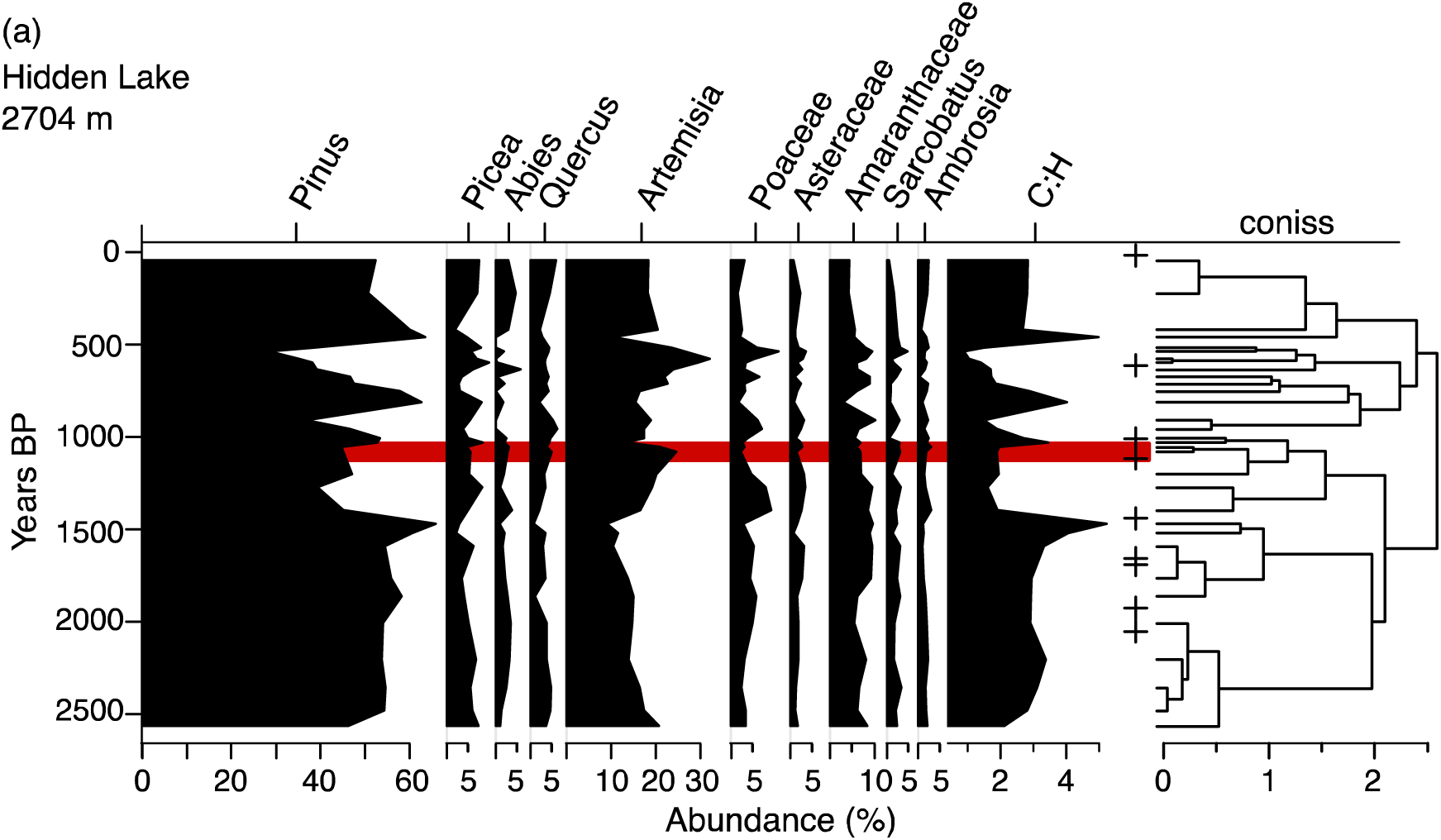
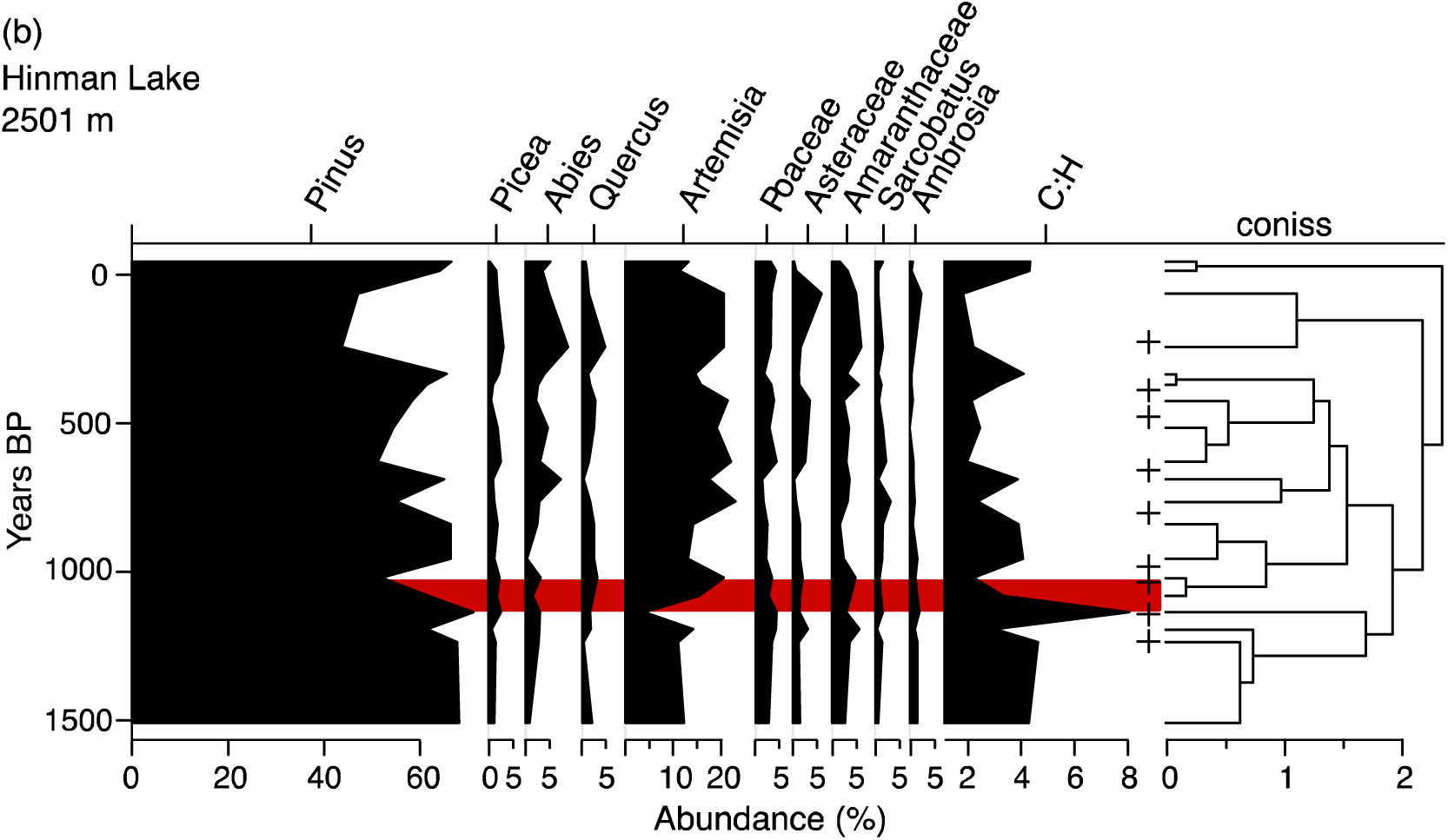
Terrestrial pollen percentages from the lowest elevation sites: Hidden Lake (A) and Hinman Lake (B). Plus symbols represent fire events detected within the individual cores and red bars highlight the century of peak landscape scale wildfires shown in Figure 7.

Finally, Hinman Lake experienced many changes like those observed at higher elevations, including maximum *Pinus* pollen percentages at 1136 BP before the local expression of the large-scale fires and a sharp rise in *Artemisia* pollen percentages to >20% (Fig. 6b). *Pinus* pollen percentages declined at the same time, ultimately reaching a minimum from ca. 300 – 100 BP when both *Abies* and Asteraceae pollen percentages also reached maxima. The C:H ratio likewise fell from >4 from ca.1500 to 1136 BP to <2 in most samples from ca. 600 – 100 BP. The decline in the interpolated C:H at 1100 BP has a 1.00 probability of a change point.

### Pollen Zones

Because of the varied local changes, a break in high order clusters of pollen samples was located between ca.1200 and 1000 BP at all sites except for Gem Lake. The boundaries between stratigraphic clusters (pollen zones), therefore, overlap in time with the most widespread fires (red bars, Figs 4 – 7). At Summit Lake, the largest separation of clusters falls between pollen samples with median ages of 1021 and 987 BP (Fig. 4b). At Seven Lakes, the second largest cluster break dates between 1275 and 1132 BP but given the age uncertainties and spacing of pollen samples, the break is not significantly different in time from the break at Summit Lake or the most widespread fires (red bar, Fig. 4). At Gold Creek Lake, a third order break falls between 1188 – 1101 BP, and at Hidden Lake between 1005 – 956 BP (Figs 5b and 6a). A high order separation between clusters also dates to 1136 – 1081 BP at Hinman Lake, although the first order break dates to ca. 300 BP (Fig. 6b). At Gem Lake, however, the largest separation between clusters dates to 780 – 705 BP (Fig. 5a), which is consistent with other high order cluster breaks at sites such as Gold Creek and Hinman that could be associated with forest changes associated with the LIA.

The median C:H ratios from the mid- and high-elevation sites (Seven, Summit, Gem, and Gold Creek lakes) summarizes the differences between the clusters of pollen samples before and after 1021 (1275 – 956) BP (Fig. 7). Before the breaks in the constrained cluster analysis (Figs 4 – 6), and the peak in the area burned, which dates to 1130 – 1030 BP (Fig. 7a; Calder et al., 2015), the median C:H ratio varied between 2.49 and 1.97. However, from 1165 – 970 BP, the median ratio declined from 2.11 to 1.67, which was the largest single period of decline since 2000 BP (Fig. 7c). Afterwards, the ratio continued to decline and varied between 1.67 and 1.36 from 970 to 60 BP, in parallel with regional cooling (Fig. 7).

**Fig. 7.**
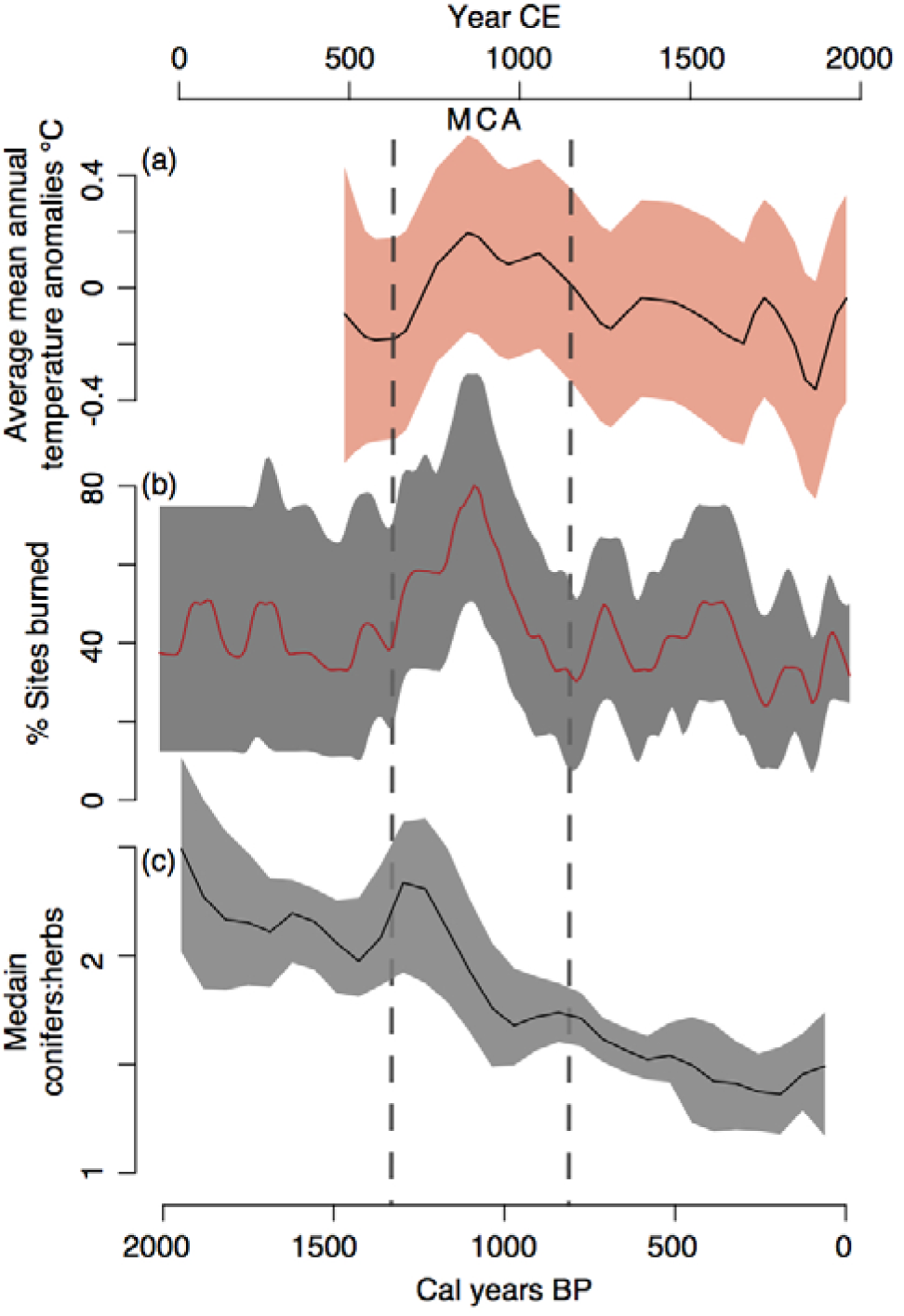
Vegetation change across the landscape after a rise in mean annual temperature in North America (a; from Trouet et al. 2013) and percent sites burned (b; from Calder et al. 2015) with median C:H ratios (c) from the four highest elevation sites (Summit Lake, Seven Lakes, Gem Lake, and Gold Creek Lake). Orange bands in (a) represent two standard errors around the mean (black), and grey bands in (b) and (c) represent 90% confidence bands around median percent sites burned and pollen ratio.

### Unconstrained cluster analysis

A cluster analysis of all the pollen samples, which was not constrained by their temporal position, indicates that the samples fall into three major groups (Fig. 8). The top cluster contains the modern sample from Summit Lake, which is the most representative of ribbon forests of any of the sites. At Summit Lake, most samples after the fires ca. 1000 years ago fall into this first, ribbon forest cluster (Fig 8). The top cluster later became important at Gem and Gold Creek lakes, although the modern samples from each of these sites, as well as early samples from Summit Lake, are grouped within the middle cluster. The middle cluster, therefore, appears to represent closed spruce-fir forests and was most important in the early part of all the records except at Hidden and Hinman lakes. The third cluster includes the low-elevation samples from Hidden and Hinman lakes, and therefore, represents mixed forests of lodgepole, aspen, spruce, and fir.

**Fig. 8.**
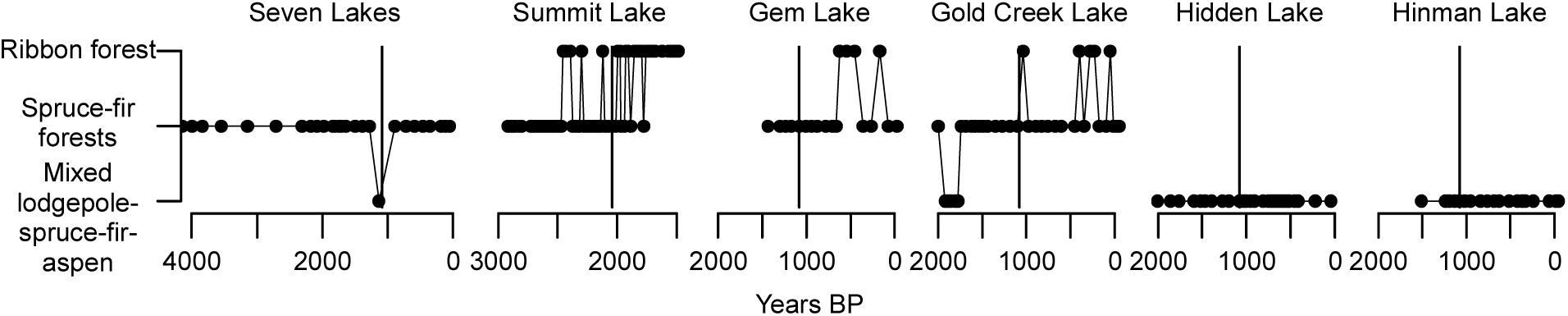
Unconstrained cluster analysis of the first three clusters from all sites separated by site through time. The vertical line indicates the time of peak landscape wildfires centered at 1130 BP.

The opening of the forests at most sites began sharply at the time of the fires at each site (Figs 4 – 7), but the cluster analysis indicates that only the vegetation changes at Summit Lake exceeded the differences between elevational zones today at that time (Fig. 8). The continued trend toward increased *Artemisia* and reduced conifer pollen percentages (e.g., Fig. 7c) was required before Gem and Gold Creek lakes shifted from one cluster to another during the LIA (Fig. 8). Thus, the ribbon forest cluster became widespread by ca. 500 BP, even though it had been rare prior to 1000 BP. Indeed, the Seven Lakes record extends to >4000 BP and Summit Lake extends to >2800 BP, but neither lake contains samples that cluster with the ribbon forests around Summit Lake today before ca. 2000 BP. Even then, the ribbon forest cluster only registers intermittently at Summit Lake from 2000 – 1000 BP, when regional isotopic datasets indicate another earlier cool or snowy interval (Anderson, 2011; Anderson, Berkelhammer, & Mast, 2015), and it only becomes important across the landscape after 1000 – 500 BP (Fig. 5). The pollen source area of Seven Lakes never became dominated by ribbon forest, but during the LIA, ribbon forests became most extensive across the Park Range based on their expansion near Gem and Gold Creek lakes (Fig. 5).

## Discussion

### Vegetation changes

The pollen data support our first hypothesis that large fires can trigger vegetation changes with legacies that persist across the landscape. Climate trends, such as cooling that culminated in the LIA after ca. 650 BP (Fig. 7a), played a first-order role in shaping forest composition (Calder and Shuman, 2017) and determined the time when the modern patterns of forest types developed. Ribbon forests, for example, became most extensive during the LIA (Fig. 8). At nearly all the study sites, however, pollen assemblages changed after the most widespread wildfires of the last 2000 years. Five of the six pollen records show high order clustering before and after the peak in the area burned by wildfires (Figs 4 – 6). The median ratio of C:H pollen indicates that extensive Medieval wildfires (Fig. 7b) accelerated a decline in tree cover (Fig. 7c) even though warming at the time (Fig. 7a) may have initially favored an expansion of forest cover (Fig. 7c), often represented by pre-fire maxima in *Pinus* pollen percentages (*Pinus* peaks below red lines in Fig. 4-5). After the fires, the relative abundance of open meadow taxa rapidly increased and remained high for the last 1000 years (Figs 4 – 7).

The vegetation changes ca. 1000 years ago also produced pollen assemblages without many earlier analogs. The cluster of pollen samples that today includes samples from the modern ribbon forest at Summit Lake only became important across the landscape in the last millennium (Fig. 8), indicating that the vegetation patterns first began to develop a modern configuration in the last 2000 years after the extensive fires ca. 1000 BP and then the cooling that marked the LIA after ca. 650 BP. Forest openings were not limited to these high elevation areas, however, because large changes were also observed near the low-elevation ecotone at Hinman Lake (Fig. 6b) and because the high-elevation ribbon forests represent only a small portion of the pollen source area of the six lakes. At Hinman Lake, forest opening may represent the expansion of meadows in areas of cold air drainage, which would also be consistent with an increase in *Abies* pollen percentages there by ca. 700 BP (Fig. 6b).

The apparent vegetation legacies of the fires persisted far longer than expected if only ecological factors, such as limitations on seed dispersal, had impacted forest recovery. The persistence of the changes may, instead, reflect the effects of regional climate changes after the wildfires, including severe Medieval droughts (Cook, Woodhouse, Eakin, Meko, & Stahle, 2004; Woodhouse, Meko, MacDonald, Stahle, & Cook, 2010) and later regional cooling or increased snow precipitation (Anderson et al., 2015; Mann et al., 2009; Trouet et al., 2013). Both drought and later cooling could have reduced post-fire recruitment (Harvey, Donato, & Turner, 2016; Peet, 1981) and prevented post-fire recovery to the previous forest state across the mountain range. As a result, the synchronized wildfires catalyzed ecological changes across the landscape, which were then sustained by long-term climate shifts favorable to open forests (Calder & Shuman, 2017; Turner, 2010). Ultimately, cooling associated with the LIA (Fig. 7a) was probably important because it could have limited germination and recruitment at high elevation and in areas of cold air drainage, such as the sagebrush park below Hinman Lake (Coop & Givnish, 2008), and ribbon forests may have been most extensive at that time (Fig. 8).

### Pre-fire conditions

Before the widespread wildfires, increases in forest cover may have created optimal conditions for the spread of large fires. As temperature began to increase at the beginning of the Medieval period (MCA in Fig. 7a), the percentages of *Pinus* or *Picea* pollen increased to a peak at 5 of the 6 sites. At Summit, Gem, Gold Creek, and Hinman lakes, *Pinus* pollen reached maxima of 38-71% at 1188 – 1136 BP. At Seven Lakes, both *Picea* and *Pinus* pollen percentages rose to near their maximum of the last 2400 years (16% and 54%, respectively). The combined records indicate that the pre-fire maximum of the median C:H ratio represents a meaningful phase of the landscape history (Fig. 7c), which could be consistent with increased forest density during the initial phases of regional warming (Fig. 7a). Regional warmth may also be indicated by peaks in the long-distance transport of *Quercus* pollen at the same time (Figs 4 and 5).

With the regional warming, conifers may have invaded high elevation meadows and areas of cold air drainage. Competitive advantages caused by the warming or outbreaks of spruce beetles (*Dendroctonus rufipennis* Kirby) could have favored *Pinus* over *Picea* at Summit Lake (Berg, David Henry, Fastie, De Volder, & Matsuoka, 2006; Veblen, Hadley, Reid, & Rebertus, 1991), which was similar to the replacement of *Picea* by *Pinus* at Gold Creek Lake during an earlier period of warmth or low snow cover ca. 2000 BP (Figs 4b and 5b; Anderson 2011; Anderson *et al.* 2015).

Peaks in *Quercus* pollen percentages at Seven and Gold Creek lakes at ca. 1000 BP may also represent a response to warming (Figs 4a and 5b). The long-distance dispersal of pollen from Gamble’s oak populations on the west slope of the Park Range could indicate that the populations expanded upslope, especially into south facing microsites, as the landscape warmed. Increased effective moisture during the last millennium may have also been a factor in their expansion (Anderson 2011; Shuman et al., 2009), while the opening of the forests around the high-elevation lakes over the last millennium may have also favored the deposition of remotely dispersed oak pollen.

### Post-fire conditions

After the widespread wildfires, the four high elevation pollen records indicate a rapid decline in conifer abundance relative to shrub and herb pollen (Fig. 7c). The open forest types that developed include the then-novel assemblage that today encompasses the high-elevation ribbon forests observed near Summit Lake (Fig. 8). Fire probably favored the rapid formation of the first large areas of ribbon forests (Calder & Shuman, 2017), and subsequent cooling favored their spread as indicated by the appearance of the same cluster at Gem and Gold Creek lakes by ca. 500 BP (Fig. 8). Increases in *Artemisia* pollen percentages at sites like Hinman Lake (Fig. 6b), however, illustrate that the forests also opened in other ways after the fires, such as through the creation of new low elevation meadows.

The recovery of *Pinus* pollen percentages after the sharp fire-related minimum at Gold Creek Lake (red bar, Fig. 5b) indicates that some successional recovery followed the fires. In fact, the response at Gold Creek Lake first includes a peak in the percentages of *Artemisia* and herbaceaous taxa at ca.1100 BP, then a brief maximum in *Pinus* pollen percentages, and finally an increase in *Picea* pollen percentages, which is not unlike the successional sequence in these forests today. The apparent successional recovery at Gold Creek Lake culminated, however, in a lower C:H ratio than observed before the wildfires (Fig. 5b). At Hidden Lake, the shift toward an open forest was not persistent and may also be consistent with its elevation, where neither high-elevation low temperatures nor winter temperature inversions in low-elevation areas would have been a factor (Fig. 6a). Also, the fires may have been less severe near Hidden Lake than in other areas.

### Why changes were detected

Detecting paleoecological changes in response to wildfire is difficult. Contemporary examples show that wildfires can trigger abrupt vegetation changes, and many paleoecological studies have attempted to understand the effects of wildfire on century to millennia time scales across a variety of biomes (Clifford & Booth, 2015; Green, 1982; Nelson, Hu, Grimm, Curry, & Slate, 2006; Umbanhowar, 2004). However, in multiple subalpine ecosystem studies, no consistent patterns were observed between detected fires and the pollen records (Minckley & Long, 2016; Shriver & Minckley, 2012). One challenge associated with detecting the influence of fires on vegetation arises from differences in pollen and charcoal source areas.

Individual lake records, like individual fire scarred trees, only record fire within a point on the landscape. As a result, charcoal peaks cannot be used to determine whether a fire burned a small or large portion of the surrounding ecosystem (Gavin, Brubaker, & Lertzman, 2003). In fact, most wildfires that produce significant charcoal accumulation peaks do not burn across the entire pollen or charcoal source area (Gavin et al. 2003), leaving vegetation intact to continue contributing pollen to a sediment record and obfuscating any record of the vegetative responses to wildfire. Analyzing pollen at one spatial and temporal scale, and charcoal at another, inevitably creates patterns related to differing scales rather than the desired wildfire-climate-vegetation interactions (Wiens 1989).

Our composite record of the percentage of sites burned per century (Fig. 7b; Calder et al., 2015) may, however, resolve the problem by sampling wildfire at spatial scales closer to the scale of the full pollen source area. Without a composite fire record it would be hard to determine *a priori* which individual lake charcoal peaks would be expected to create a large change in the pollen records. Most local fires (plus symbols, Figs 4 – 6) did not produce any apparent changes in our pollen records. Localized fire events at Hidden Lake ca. 650 and ca.1450 BP may be notable exceptions because they were followed by changes in the pollen assemblages consistent with an opening of the surrounding landscape, but this type of response in these individual records is the exception rather than the rule. The coincidence of widespread evidence of fires (Fig. 7b) and sharp, widespread declines in tree abundances at ca. 1100 BP (Figs 4 – 6 and 7c), however, are consistent with the expectation that the spread of exceptionally large fires could substantially influence the mix of pollen dispersed across the landscape. That the change persisted because of subsequent climatic limitations on forest recovery further increased the likelihood of detection.

### Fire feedbacks

The increase in non-arboreal pollen percentages at the high elevations lakes likely represents the development of ribbon forests, large meadows, and potentially new areas of tundra across the landscape (Fig. 7c). The opening of the high elevation forests could have created fuel breaks across the mountain range, which would be consistent with our second hypothesis that vegetation changes could feedback to alter fire regimes. Such limits on the spread of wildfire may explain why the area burned per century declined after ca. 1100 BP even while warmth persisted in the region (Fig. 7a, b), especially if severe Medieval droughts delayed forest recovery (Calder et al., 2015; Cook et al., 2004; Woodhouse et al., 2010). Indeed, isolated high-elevation sites, like Seven Lakes, recorded almost no fires in the past millennium once the pollen records show an increase in non-arboreal pollen (Fig. 4a; Calder et al. 2014). The changes in wildfire and vegetation indicate that the relationships among components of the fire-regime triangle (Fig. 1) were sensitive to small (~0.5 °C) climate changes and the particular sequence of events.

The coincident decline in arboreal pollen percentages and in wildfire frequency when temperatures remained elevated (ca. 1000 BP in Fig. 7) indicates an important vegetation feedback in the fire-regime triangle (Fig. 1d). The change supports our first hypothesis that fires in conifer forests can facilitate critical transitions and the development of multiple states, defined by high and low forest cover that depends upon the fire history (Scheffer, Hirota, Holmgren, Van Nes, & Chapin, 2012). Open areas may then further limit the spread of fire. In boreal forests, past fire-vegetation feedbacks increased less flammable deciduous taxa and, thus, limited the area burned by subsequent fires even after regional temperatures increased (Girardin et al., 2013; Kelly et al., 2013; J. A. Lynch, Clark, Bigelow, Edwards, & Finney, 2002). Likewise, negative fire-vegetation feedbacks indicate that models based strictly on climate-fire relationships will over-predict the area burned in boreal forests (Heon, Arseneault, & Parisien, 2014). Future climate change will likely synchronize widespread fires across subalpine landscapes (Westerling, Turner, Smithwick, Romme, & Ryan, 2011), but the evidence here indicates that climate-fire-vegetation feedbacks could alter forest density or composition at high elevations, and limit or modify the long-term wildfire responses (Fig. 7).

## Conclusions

Climate change influences both vegetation and wildfire (Fig. 1), and the history of the Mount Zirkel Wilderness indicates that small temperature increases (~0.5 °C) can result in widespread wildfires and large vegetation changes. Wildfire regimes responded nonlinearly to temperature as the percentage of sites burned declined while temperatures were still elevated (Calder et al., 2015). The nonlinearity most likely arose because vegetation also responded nonlinearly through interacting effects of wildfire and climate change. Fires killed trees across the landscape, and the sharp shift in vegetation patterns and composition altered fuel structures and continuity. The vegetation change, thus, prevented additional large fires from burning, and as a result, the ecosystem, including both its vegetation and disturbance regimes, shifted abruptly. Novel patterns and forest types emerged, especially as climate continued to change and shape the legacies of the fires (Fig. 8). These changes underscore the potential for forests to undergo critical transitions as ongoing climate changes alter the relationships among climate, vegetation, and wildfire.

## Author Contributions

W.J.C. and I.V. generated the data and W.J.C. and B.S. developed the study design, extracted the sediment cores, analyzed the data, and wrote the manuscript. All authors contributed to the drafts and gave final approve for publication.

### Acknowledgements

We thank Jeremiah Marsicek, Grace Carter, Dusty Parker, Cody Stopka, Keith Ingledew, Devin Hougardy, Corianne Calder, Rhowe Stefanski, Ethan Harris, Ashley Harris, Marc Serravezza, Aaron Condie, Danny Vecchio, and David Webster for help coring and with lab work. The National Science Foundation Grant BCS-0845129 (to B.N.S.) supported this project.

## Data Accessibility

Charcoal data for the wildfire reconstructions (Calder et al., 2015) are available through the International Multiproxy Paleofire Database (www.ncdc.noaa.gov/data-access/paleoclimatology-data/datasets/fire-history). The pollen data will be made available upon publication in the Neotoma Paleoecology Database (www.neotomadb.org/).

